# A library-on-library screen reveals the breadth expansion landscape of a broadly neutralizing betacoronavirus antibody

**DOI:** 10.1101/2024.06.06.597810

**Authors:** Marya Y. Ornelas, Wenhao O. Ouyang, Nicholas C. Wu

## Abstract

Broadly neutralizing antibodies (bnAbs) typically evolve cross-reactivity breadth through acquiring somatic hypermutations. While evolution of breadth requires improvement of binding to multiple antigenic variants, most experimental evolution platforms select against only one antigenic variant at a time. In this study, a yeast display library-on-library approach was applied to delineate the affinity maturation of a betacoronavirus bnAb, S2P6, against 27 spike stem helix peptides in a single experiment. Our results revealed that the binding affinity landscape of S2P6 varies among different stem helix peptides. However, somatic hypermutations that confer general improvement in binding affinity across different stem helix peptides could also be identified. We further showed that a key somatic hypermutation for breadth expansion involves long-range interaction. Overall, our work not only provides a proof-of-concept for using a library-on-library approach to analyze the evolution of antibody breadth, but also has important implications for the development of broadly protective vaccines.

## INTRODUCTION

Many RNA viruses have high genetic diversity and undergo rapid antigenic drift due to the selection pressure from host humoral immune responses. As a result, it is often challenging to develop effective vaccines with a high breadth of protection against RNA viruses. Nevertheless, in the past two decades, a number of broadly neutralizing antibodies (bnAbs) against different RNA viruses, including influenza virus^1–4^, human immunodeficiency virus^5–7^, coronavirus^8–11^, and flavivirus^12,13^, have been isolated. These bnAbs protect against antigenically distinct strains of a given virus species, or even genus and family. The discovery of bnAbs has provided crucial insights into the development of broadly protective vaccines^9,14–16^.

Most bnAbs evolve from an unmutated common ancestor (UCA) with narrow binding specificities^17–20^. Their cross-reactivity breadth is subsequently acquired through the accumulation of somatic hypermutations (SHMs) during affinity maturation^21–23^. Yeast display is a common approach for studying the evolutionary landscapes of antibody affinity maturation^24–32^. This process typically involves the construction of an antibody mutant library, which is then displayed on yeast surface and selected for antigen binding^24,26–28,30–32^. However, conventional yeast display selection focuses on only one antigen at a time, which imposes a challenge in studying the evolution of antibody breadth. Improving binding affinity of an antibody against one antigenic variant sometimes diminishes its binding affinity against another variant^27^. Yet, evolution of breadth requires improvement of binding affinity across multiple antigenic variants simultaneously. As a result, characterization of the evolutionary pathways for antibody breadth expansion requires selection against multiple antigenic variants in parallel.

In the past few years, studies of human antibody responses to SARS-CoV-2 have led to the discovery of bnAbs that target the highly conserved stem helix peptide in the S2 domain of the coronavirus spike glycoprotein^8,8,33,34^. S2P6, which was isolated from a COVID-19 convalescent individual, is a representative bnAb to stem helix peptide^10^. While S2P6 neutralizes antigenically distinct betacoronavirus (β-CoV) strains including SARS-CoV-2, antibody binding data suggest that it arose in response to HCoV-OC43 infection and gained cross-reactivity to other β-CoV strains via SHMs^10^. Given that stem helix peptide is a target for the development of broadly protective coronavirus vaccines^35^, it is important to understand the evolutionary trajectories that lead to breadth expansion of bnAbs to the stem helix peptide.

Recently, a yeast display platform for coevolving protein-protein interfaces was developed^36^. Here, we adopted this platform to screen a library of 27 unique β-CoV stem helix peptides against a mutant library of S2P6 encoding all combinations of SHMs that lie on or near the paratope. This approach enabled us to map the binding affinity landscapes of S2P6 against stem helix peptides from all β-CoV subgenera. We observed weak correlations of binding affinity landscapes of S2P6 across different β-CoV stem helix peptides, indicating that the effect of a given SHM on S2P6 binding could vary depending on the sequence of the target stem helix peptide. At the same time, several key SHMs for breadth expansion could be identified and experimentally validated. Our results further highlight the importance of long-range interaction in the affinity maturation of S2P6. Throughout this study, the Kabat numbering scheme is used for antibody residues unless otherwise stated.

## RESULTS

### Detecting interaction between S2P6 and SARS-CoV-2 stem helix peptide by yeast display

To determine if we could adopt a previously developed protein-protein coevolution platform^36^ to screen an antibody mutant library against an antigen library, a pilot experiment was performed using the SARS-CoV-2 stem helix peptide and S2P6^10^. We cloned a yeast display construct that consisted of (from N-terminal to C-terminal): Aga2p, stem helix peptide (SP), 3C protease cleavage site and linker (3C), S2P6 in single-chain variable fragment (scFv) format, and HA tag (**Fig. 1A and S1**). After the construct was displayed on the yeast surface, the yeast cells were treated with 3C protease. If S2P6 bound to the stem helix peptide, the HA tag would be retained at the yeast cell surface. If S2P6 did not bind to the stem helix peptide, the HA tag would be lost (**Fig. 1A**). For the yeast displaying the SARS-CoV-2 stem helix peptide and S2P6, the HA tag was readily detected both before and after protease treatment (**Fig. 1D**). This result indicated that S2P6 bound strongly to the SARS-CoV-2 stem helix peptide on yeast surface as expected. By contrast, for the yeast displaying the SARS-CoV-2 stem helix peptide and S2P6 UCA, which was the fully germline-reverted S2P6^10^, we observed loss of the HA tag after protease treatment (**Fig. 1D**). This observation indicated that the S2P6 UCA did not bind to the SARS-CoV-2 stem helix peptide, consistent with the previous study^10^. Overall, our pilot experiment demonstrated that the interaction between S2P6 and stem helix peptide could be captured by the yeast display platform that was previously developed for studying protein-protein coevolution^36^.

**Figure 1.**
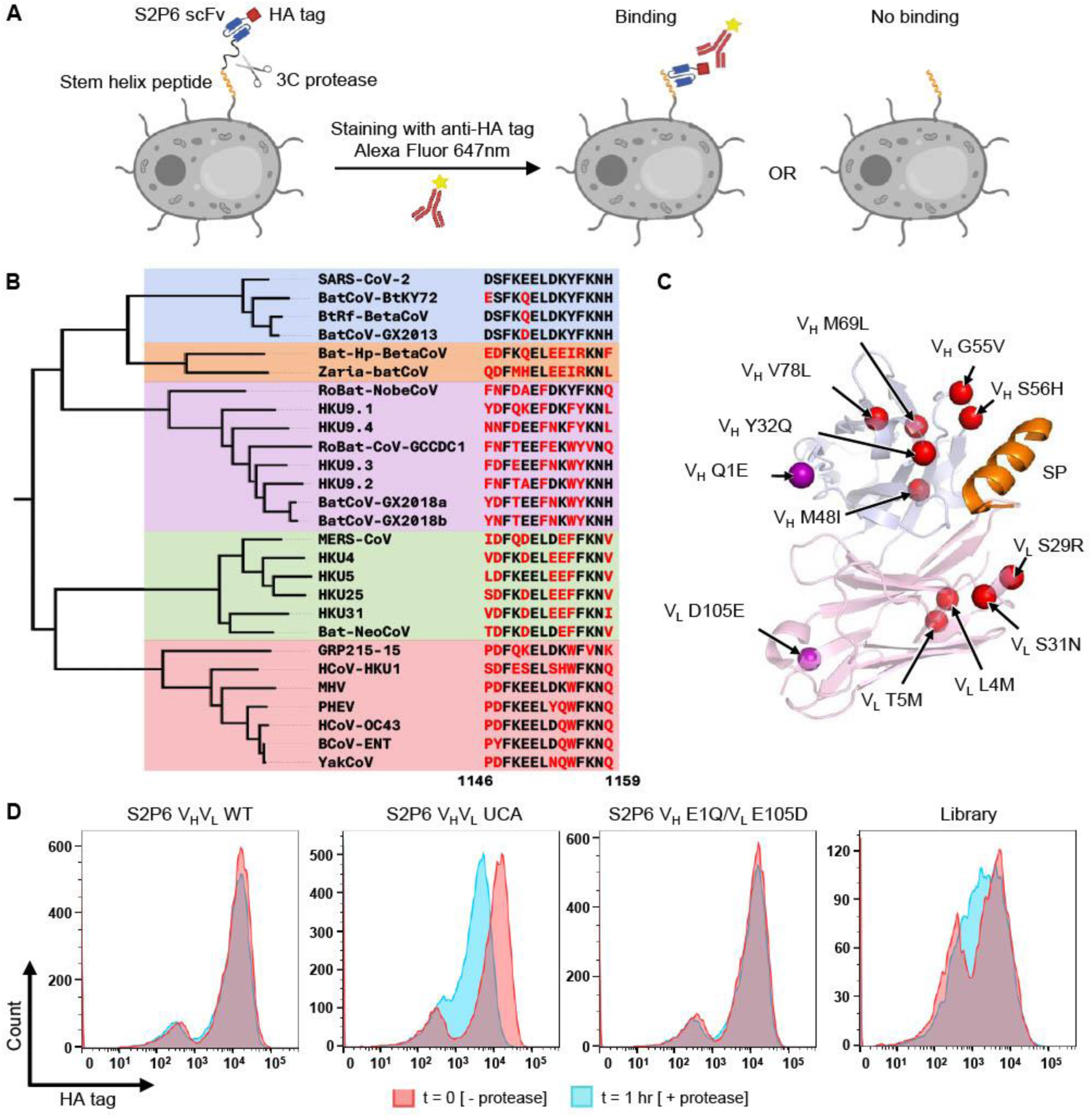
Overview of library-on-library experimental design. **(A)** Each yeast cell displayed a S2P6 variant in single-chain variable fragment (scFv) format and a stem helix peptide variant with a 3C protease cleavage site in between. After treatment with 3C protease and staining with anti-HA tag Alexa Flour 647 nm, S2P6-SP binding could be assessed using flow cytometry. **(B)** A phylogenetic tree was constructed using the spike sequences of the β-CoV strains included in this study. Sequences of their spike stem helix peptides (residues 1146-1159, SARS-CoV-2 numbering) are shown on the right. Box color denotes β-CoV subgenus: sarbecovirus (blue), hibecovirus (orange), nobecovirus (purple), merbecovirus (green), and embecovirus (red). **(C)** The structure of S2P6 Fab in complex with SARS-CoV-2 stem helix peptide (orange) is shown (PDB 7RNJ)^10^, with heavy chain variable domain (V_H_) in light blue and light chain variable domain (V_L_) in light pink. Somatic hypermutations included in our library design are shown in red and those excluded are in purple. **(D)** Flow cytometry was performed to analyze the HA tag signal (Alexa Flour 647 nm) of yeast cells displaying S2P6 WT, S2P6 UCA, and S2P6 V_H_ E1Q/V_L_ E105D with SARS-CoV-2 stem helix peptide, as well as the combination variant library. “- protease” and “+ protease” indicate before and after protease treatment, respectively.

### Screening an S2P6 mutant library against a stem helix peptide library

We sought to analyze the affinity maturation pathway of S2P6 by screening an S2P6 mutant library encoding all possible combinations of SHMs against a library of 27 unique β-CoV stem helix peptides (**Fig. 1B and Table S1**). Among the 12 SHMs in S2P6 (**Fig. 1C and S2A-B**), V_H_ Q1E and V_L_ D105E were distal from the binding interface. Reverting these positions in S2P6 to the germline sequence (V_H_ Q1 and V_L_ D105) did not result in loss of HA tag in our yeast display system after protease cleavage (**Fig. 1D**). This observation indicated that V_H_ Q1E and V_L_ D105E played a minimal, if any, role in the affinity maturation of S2P6. As a result, we moved forward with constructing the S2P6 mutant library without V_L_ D105E and V_H_ Q1E mutations (**Fig. S2C**). In other words, our S2P6 mutant library contained 1,024 variants (2^10^ = 1,024). Together with the 27 unique β-CoV stem helix peptides and one negative control peptide, there were a total of 28 peptide variants × 1,024 S2P6 variants = 28,672 combination variants in our library-on-library screen.

To track the frequency of different combination variants in our library, next-generation sequencing was required. While the region of interest in the previous study that developed the protein-protein coevolution platform was <600 bp^36^, which is compatible with Illumina short-read sequencing, the amplicon spanning both S2P6 scFv and stem helix peptide was too long (>800 bp). To circumvent this issue, we adopted a barcoding strategy from a previous study^37^. Briefly, we included a 16-nucleotide “barcode” downstream of the coding region, such that different combination variants had their own barcodes (**Fig. S1 and S3**). The linkages between individual combination variants and their corresponding barcodes were then mapped by PacBio sequencing, which had sufficiently long read lengths to cover both S2P6 scFv and stem helix peptide but insufficient read depths to track the frequency of each combination variant in the library. Subsequently, the frequency of different variants could be tracked by sequencing the barcode region using Illumina short-read sequencing.

Flow cytometry analysis showed that combination variants in our library had a wide range of binding affinity (**Fig. 1D**). The library was subjected to fluorescence-activated cell sorting both before and after protease cleavage (**Fig. S4**). Four bins were collected from each sort according to the amount of HA tag detected on the yeast surface. The frequency of each combination variant in each bin was quantified by next-generation sequencing as described above. An expression score and a binding score were then computed for each combination variant based on its frequency distribution in different bins, hence a proxy for HA tag signal, before and after protease cleavage, respectively (**Table S2, see Materials and Methods**). As shown previously, binding affinity strongly correlates with HA tag signal after protease cleavage^36^. Combination variants in our library had a broad distribution of binding scores (**Fig. S5A**), consistent with the flow cytometry analysis of the library (**Fig. 1D**). At the same time, they had a narrow distribution of expression scores (**Fig. S5B**), indicating that most combination variants expressed well. We also observed a Pearson correlation of 0.70 between the binding scores of two independent experimental replicates (**Fig. S5C**), demonstrating reproducibility of our library-on-library screen. By contrast, the Pearson correlation between binding score and expression score was 0.29, showing that the expression levels of combination variants had a relatively small influence on their binding scores (**Fig. S5D**).

### Breadth expansion of S2P6 is restricted by sequence variations of stem helix peptides

To analyze the breadth expansion of S2P6, we computed the average binding scores of the S2P6 mutant library for each β-CoV stem helix peptide (**Fig. 2A**). High average binding scores were observed for stem helix peptides from all sarbecovirus strains as well as embecovirus strains, except PHEV and HCoV-HKU1. Nevertheless, the average binding scores for stem helix peptides from PHEV and HCoV-HKU1 were still higher than the negative control peptide. For stem helix peptides from the embecovirus, amino acid variants at position 1153 appeared to play a critical role in binding to S2P6. At position 1153, YakCoV, HCoV-HKU1, and PHEV have N1153, S1153, and Y1153, respectively, whereas all other embecovirus strains have D1153 **(Fig. 1B)**. As previously shown by a mutational analysis of SARS-CoV-2 stem helix peptide, D1153 has the highest binding activity to S2P6, followed by N1153, S1153, and then Y1153^10^. Consistently, stem helix peptides from those embecovirus strains with D1153 had the highest average binding scores in our data, followed by YakCoV (N1153), HCoV-HKU1 (S1153), then PHEV (Y1153).

**Figure 2.**
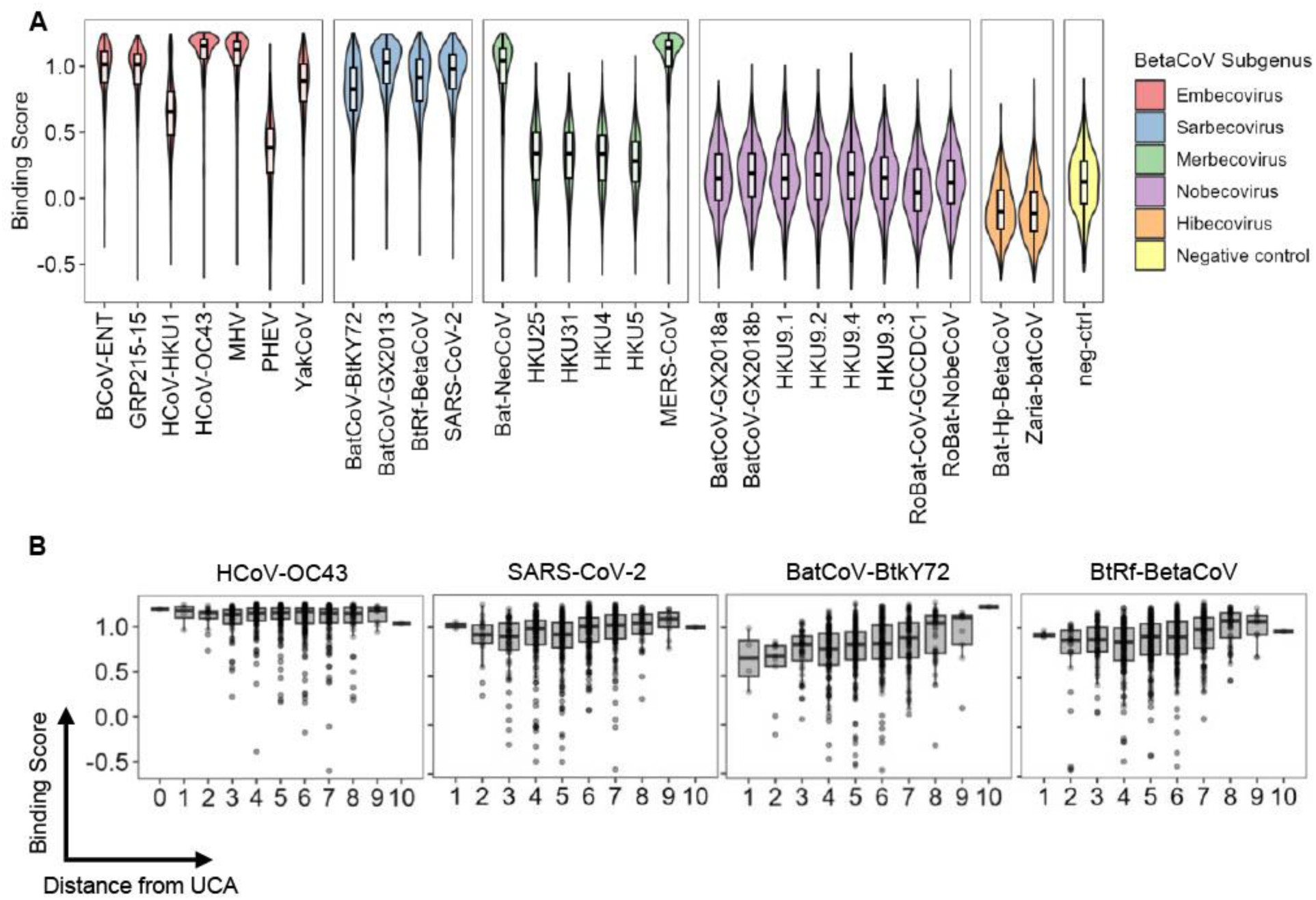
Analysis of S2P6 binding landscapes across stem helix peptides from all β-CoV subgenus. **(A)** Average binding scores of S2P6 variants to the indicated stem helix peptide variants are shown as a violin plot and a boxplot. SARS-CoV-2 stem helix peptide with two alanine mutations introduced (DSAKEALDKYFKNH), which has previously been shown to abolish binding to S2P6^10^, was used as a negative control. **(B)** Binding scores of S2P6 variants with different numbers of somatic hypermutations (i.e. distance from UCA) to HCoV-OC43, SARS-CoV-2, BatCoV-BtkY72, and BtRf-BetaCoV stem helix peptides are shown. Each data point represents the one S2P6 variant. For the boxplot, the middle horizontal line represents the median. The lower and upper hinges represent the first and third quartiles, respectively. The upper whisker extends to the highest data point within a 1.5× inter-quartile range (IQR) of the third quartile, whereas the lower whisker extends to the lowest data point within a 1.5× IQR of the first quartile.

Low average binding scores were observed for stem helix peptides from all merbecovirus strains, except MERS-CoV and Bat-NeoCoV. Notably, both MERS-CoV and Bat-NeoCoV have D1153 whereas all other merbecovirus strains have E1153 (**Fig. 1B**), again substantiating the importance of Asp at this position for S2P6 binding^10^. All hibecovirus strains also have E1153, which explained the low average binding scores to their stem helix peptides. Furthermore, stem helix peptides from all nobecovirus strains had low average binding scores. While the stem helix peptides from other β-CoV strains have L1152, all nobecovirus strains have F1152 (**Fig. 1B**), which has previously been shown to abolish binding to S2P6^10^.

### S2P6 binding landscape varies across stem helix peptides

It is known that S2P6 UCA binds strongly to the stem helix peptides of HCoV-OC43, but not those from the other two β-CoV strains that circulate in human population, namely HCoV-HKU1 and SARS-CoV-2^10^. As a result, S2P6 is hypothesized to have been initially elicited in response to HCoV-OC43 infection with specificity broadened through subsequent recall response to either or both HCoV-HKU1 and SARS-CoV-2 infections^10^. Consistently, even S2P6 variants with a low number of SHMs had high binding scores for HCoV-OC43 stem helix peptide (**Fig. 2B**). By contrast, the binding scores for stem helix peptides from all sarbecovirus strains, namely SARS-CoV-2, BatCoV-BtkY72, BtRf-BetaCoV, and BatCoV-GX2013, increased as S2P6 accumulated more SHMs (**Fig. 2B and S6**).

We further aimed to compare the binding landscapes of S2P6 across different β-CoV stem helix peptides (**Fig. 3 and S7**). Weak correlations of binding scores were observed among stem helix peptides from sarbecovirus strains (Pearson correlation = 0.14 to 0.21, **Fig. 3**). There was also a weak correlation of binding scores between BCoV-ENT (embecovirus) and sarbecovirus stem helix peptides (Pearson correlation = 0.12 to 0.26). In comparison, the correlations of binding scores between experimental replicates for each of these stem helix peptides were higher (Pearson correlation = 0.35 to 0.41, **Fig. 3**). Similarly, there was a lack of correlation of binding scores among embecovirus and merbecovirus stem helix peptides (Pearson correlation = -0.13 to 0.08), whereas moderate correlations of binding scores were observed between their experimental replicates (Pearson correlation = 0.34 to 0.56). These results demonstrated that the S2P6 binding landscape differed among stem helix peptides from different β-CoV strains. In other words, depending on the sequence of the target stem helix peptide, a given SHM of S2P6 could have different effects on binding affinity.

**Figure 3.**
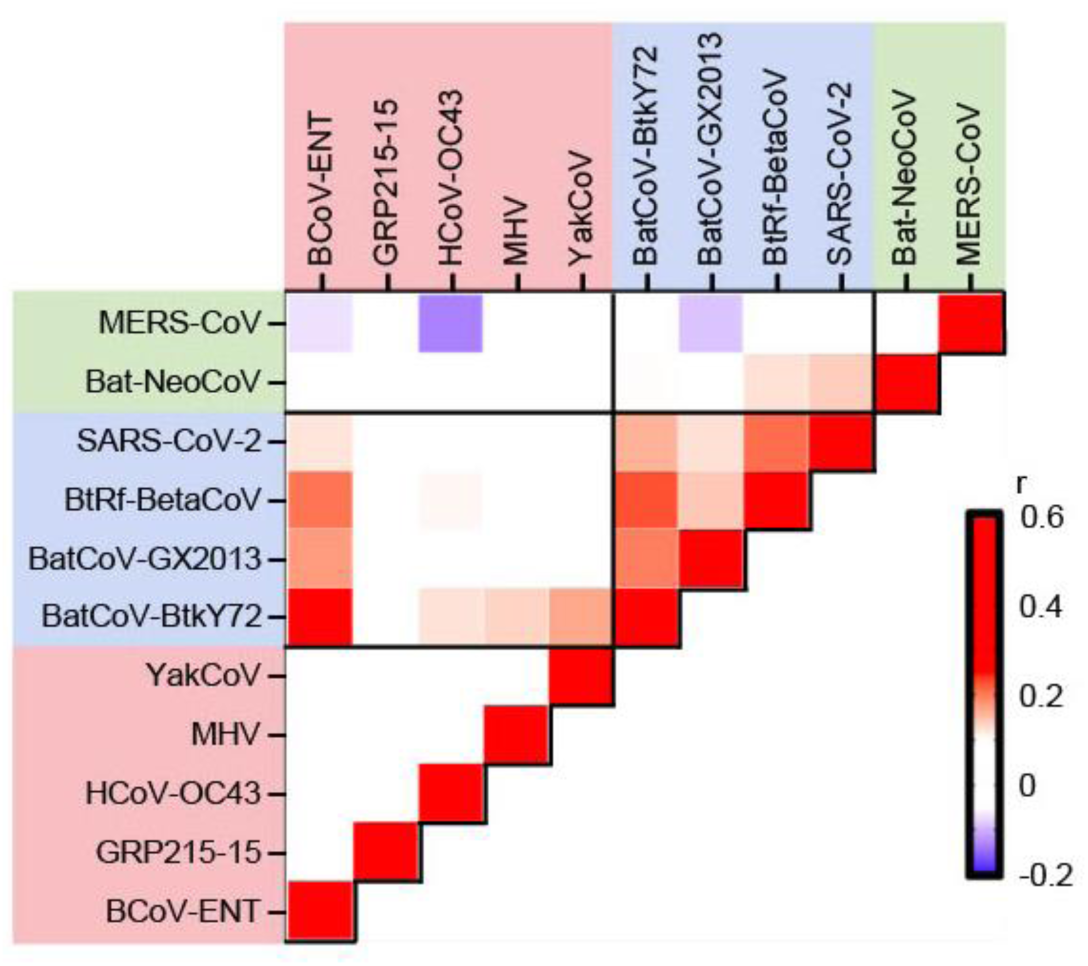
Correlation of S2P6 binding landscapes between stem helix peptides. Pairwise Pearson correlations of binding scores between stem helix peptides are shown. Only those stem helix peptides with an average binding score >0.75 are analyzed here. Red indicates positive correlation coefficient, whereas blue indicates negative correlation coefficient. Diagonal represents the Pearson correlation coefficients between experimental replicates of the indicated stem helix peptides. Shading of axis labels indicates β-CoV subgenus: merbecovirus (green), sarbecovirus (blue), and embecovirus (red).

### Affinity maturation of S2P6 involves long range interaction

Next, we aimed to analyze the impacts of individual SHMs on S2P6 binding landscape. Briefly, the S2P6 binding landscape was decomposed into additive effects of individual SHMs and pairwise epistasis effects between SHMs (**Fig. 4A and S8, see Materials and Methods**)^38^. Three SHMs on heavy chain, namely V_H_ Y32Q, V_H_ M48I, and V_H_ S56H, stood out as having positive additive effects on binding to stem helix peptides from most β-CoV strains (**Fig. 4A**). To experimentally validate this finding, these mutations were introduced individually and in combinations (single, double, and triple mutants) to S2P6 UCA and expressed as fragment antigen-bindings (Fabs). Biolayer interferometry experiments showed that each single mutation improved the binding response to the stem helix peptides from BatCoV-BtkY72, SARS-CoV-2, BtRf-BetaCoV, and MHV in comparison to S2P6 UCA (**Fig. 4B and S9**). Consistently, reverting V_H_ Q32 to V_H_ Y32 has been shown to decrease the binding affinity of S2P6 to multiple β-CoV stem helix peptides^10^. Nevertheless, among the three SHMs tested, V_H_ M48I conferred the largest improvement in binding response to all four tested stem helix peptides. V_H_ M48I alone led to a higher binding response than V_H_ Y32Q/S56H and V_H_ Y32Q/M48I double mutants for BatCoV-BtkY72, SARS-CoV-2, and BtRf-BetaCoV stem helix peptides. Similarly, for the stem helix peptides from BatCoV-BtkY72, SARS-CoV-2, and BtRf-BetaCoV, V_H_ M48I/S56H double mutant had higher binding response than the V_H_ Y32Q/M48I/S56H triple mutant. Since V_H_ Y32Q did not improve the binding response in the presence of V_H_ M48I, our data also suggested that negative epistasis existed between V_H_ Y32Q and V_H_ M48I, which was also observed in our decomposition analysis, albeit mildly (**Fig. S8**).

**Figure 4.**
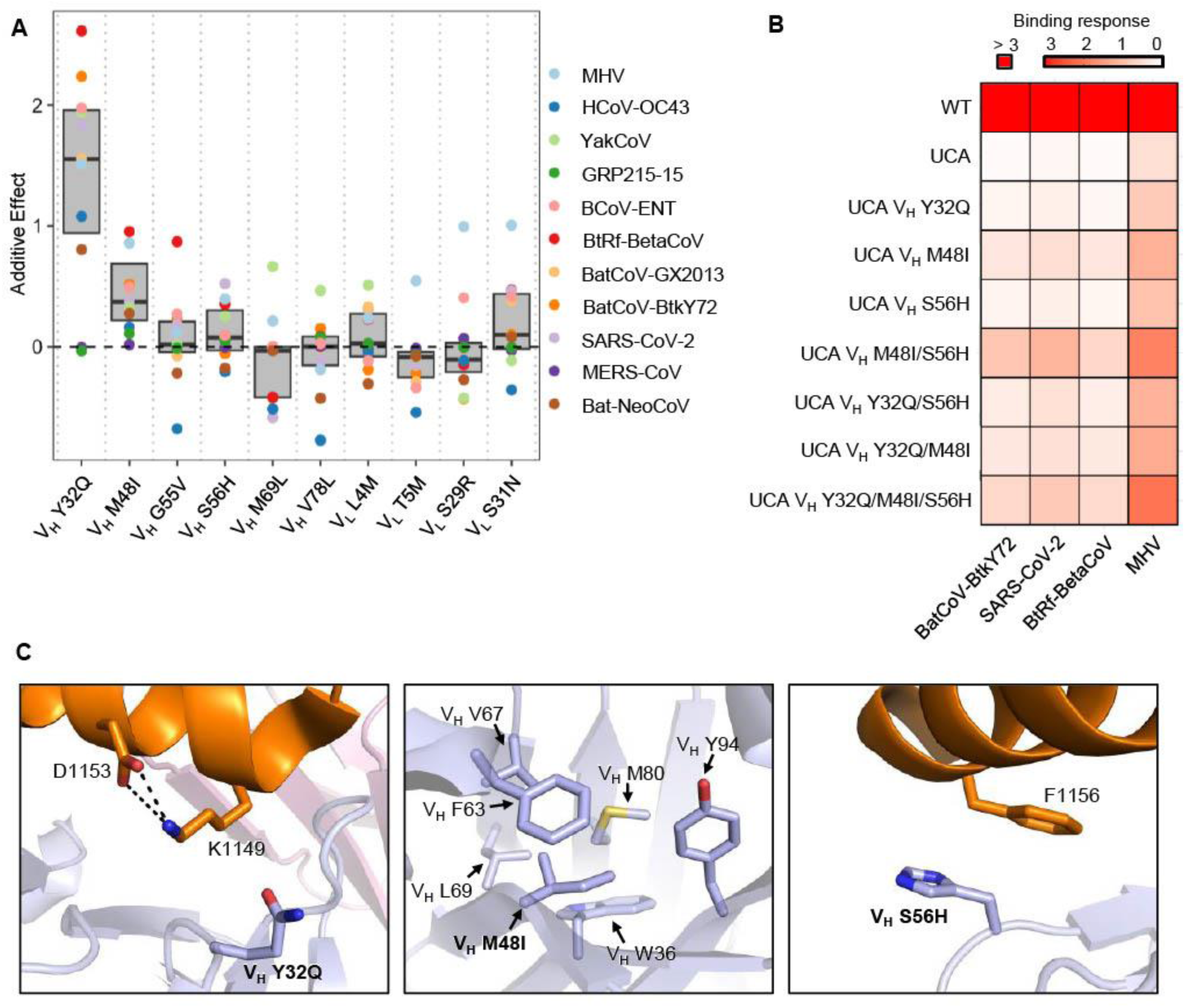
Key somatic hypermutations for breadth expansion. **(A)** The additive effects of the indicated somatic hypermutations on binding to different stem helix peptides are shown. Only those stem helix peptides with an average binding score >0.75 are included here. **(B)** Response levels (nm) of S2P6 WT, UCA, and UCA mutant Fabs binding to BatCoV-BtkY72, SARS-CoV-2, BtRf-BetaCoV, and MHV stem helix peptides were measured using biolayer interferometry. **(C)** Structural analysis of somatic hypermutations V_H_ Y32Q, V_H_ M48I, and V_H_ S56H is shown. SARS-CoV-2 stem helix peptide and S2P6 V_H_ are shown in orange and blue, respectively (PDB: 7RNJ)^10^.

Since V_H_ Y32Q and V_H_ S56H are in the paratope of S2P6^10^ (**Fig. 4C**), it may not be surprising that they could improve the binding of S2P6 to β-CoV stem helix peptides. By contrast, V_H_ M48I is not in the paratope and instead resides in the hydrophobic core of the heavy chain variable domain (**Fig. 4C**). This observation indicated that V_H_ M48I improved the interaction between S2P6 and β-CoV stem helix peptides via long-range interaction. In fact, it is not uncommon for non-paratope SHMs to improve antibody binding affinity by modulating the conformation of complementarity determining region (CDR) loops that are part of the paratope^17,39^. It is possible that a similar mechanism applies to V_H_ M48I in S2P6, although additional structural analysis is required to verify this speculation.

## DISCUSSION

Conventional yeast display experiments enable high-throughput analysis of antibody affinity maturation pathways^28,30,31,40^. However, they are often limited to analyzing antibody binding to one target antigen at a time, which restricts our understanding in the evolution of cross-reactivity breadth. A platform that allows analysis of affinity maturation pathways to multiple antigenic variants in parallel is therefore needed. In this study, we addressed this gap by determining the binding affinity landscapes of S2P6^10^ against 27 unique stem helix peptides across all β-CoV subgenera in a single yeast display experiment. Our work provides not only a proof-of-concept for using a yeast display library-on-library approach to analyze the evolution of antibody cross-reactivity breadth, but also molecular insights into the development of broadly protective coronavirus vaccines.

A notable finding in this study is the weak correlations of S2P6 binding landscapes among different β-CoV stem helix peptides. This indicates the varying effects of a given SHM on binding affinity to different β-CoV stem helix peptides. Consistently, our decomposition analysis showed that most SHMs had varying additive effects. Similar observations have also been made with an influenza bnAb in a previous study^30^. Nevertheless, we also identified SHMs that improve the binding of S2P6 to most β-CoV stem helix peptides. An effective broadly protective vaccine should facilitate the acquisition of SHMs that improve binding affinity to many antigenic variants. On the other hand, SHMs that confer trade-offs for binding different antigenic variants should be avoided. Consequently, future studies need to focus on the structural mechanisms underlying the differential impacts of SHMs on breadth expansion, which in turn will be informative for the development of vaccines that optimally elicit bnAbs.

Using library-on-library approaches to screen antibody-antigen interactions has been described since at least 15 years ago, based on co-selection of a phage-displayed antibody library and a yeast-displayed antigen library^41^. Since this strategy requires cell sorting into 96-well plates followed by sequencing the antibody and antigen in each well individually^41^, its throughput is low. More recently, library-on-library approaches that utilize lentivirus-displayed antigen library and human B cell library have been described^42,43^. However, the throughput of these approaches remains moderate due to their reliance on single-cell sequencing, which analyzes at most ∼10^4^ cells per sample. By contrast, the yeast display library-on-library approach in our present study, which was adopted from Yang et al.^36^, has a much higher throughput since it is compatible with amplicon-based next-generation sequencing. If nucleotide barcode is needed as described in this study, the maximum throughput would be ∼10^6^, as determined by the throughput of a PacBio sequencing run. In fact, another yeast display library-on-library approach has been described previously that relies on yeast mating and is also compatible with amplicon-based next-generation sequencing^44^. Nevertheless, this approach requires an engineered yeast strain that is not commercially available^44^. In comparison, our approach is more accessible since it largely follows conventional yeast display protocol without the need of any special reagents.

We acknowledge that our proof-of-concept in this study is based on a small peptide antigen. However, given that larger antigens, such as influenza hemagglutinin, HIV gp120, and SARS-CoV-2 receptor-binding domain, have been successfully displayed on the yeast surface^45–47^, we anticipate that our library-on-library approach can be applied to a wide range of antibody-antigen pairs. In addition, since our approach does not rely on yeast biology, unlike the yeast mating-based approach mentioned above^44^, it can potentially be adopted to a mammalian cell display system, which is essential for even larger antigens or if glycosylation matters. Future studies should continue to improve existing library-on-library approaches for screening antibody-antigen interactions as well as develop new ones.

## MATERIALS AND METHODS

### Yeast display plasmid

A DNA fragment encoding (from N-terminal to C-terminal) a cMyc tag, SARS-CoV-2 spike stem helix peptide sequence (residues 1146-1159), 3C protease cleavage site and linker, S2P6 heavy chain variable domain (V_H_) and light chain variable domain (V_L_) connected with a GS flexible linker, and an HA tag was synthesized as an eBlock (Integrated DNA Technologies). Subsequently, this fragment was cloned into the pCTcon2 vector (Addgene, Cat. No. 41843)^48^ in frame with the Aga2 coding region at the N-terminal end using Gibson assembly (**Fig. S1**).

### Construction of the combination variant library

The cMyc-SP-3C-S2P6_VH-GS-S2P6_VL-HA library insert was divided into six fragments and assembled using overlap PCR (**Fig. S3**). All fragments were gel-purified using a Monarch DNA Gel Extraction Kit (NEB). To generate the full-length library insert, all fragments were pooled together and underwent 10 cycles of PCR with the following conditions: 98 °C for 30 s, 10 cycles of (98 °C for 10 s, 50 °C for 15 s, 72 °C for 15 s), 72 °C for 2 min, 4 °C indefinitely. Extension primers Fragment 1_F and Fragment 6_R_3 were added into the reaction, which were then ran for another 25 cycles using conditions: 98 °C for 30 s, 25 cycles of (98 °C for 10 s, 55 °C for 5 s, 72 °C for 15 s), 72 °C for 2 min, 4 °C indefinitely. The full-length library insert was gel-purified. To generate the linearized vector for the library, the plasmid encoding the SARS-CoV-2 stem helix peptide and S2P6 WT, which was described above, was used as a template for PCR using primers S2P6_vector_F and S2P6_vector_R. The linearized vector was digested with DpnI (NEB) for 2 h at 37 °C and gel purified. All PCRs were performed using PrimeSTAR Max polymerase (Takara Bio). The sequences of all primers in this study can be found in **Table S3**. All primers in this study were synthesized by Integrated DNA Technologies.

### Yeast transformation

Saccharomyces cerevisiae EBY100 cells (American Type Culture Collection, Cat. No. MYA-4941) were grown overnight in YPD medium (1% w/v yeast nitrogen base, 2% w/v peptone, 2% w/v D(+)-glucose) at 30 °C with shaking at 225 rpm until OD_600_ reached 3. An aliquot of the overnight culture was used to inoculate 100 mL fresh YPD media with an initial OD_600_ of 0.3. This culture was incubated at 30 °C with shaking at 225 rpm until OD_600_ reached 1.6. The yeast cells were collected by centrifugation at 4600 × g for 3 min at room temperature. YPD media was removed, and the cell pellet was washed twice with 50 mL ice-cold water, and then once with 50 mL of ice-cold electroporation buffer (1 M sorbitol and 1 mM calcium chloride). The cells were then resuspended in 20 mL conditioning media (0.1 M lithium acetate and 10 mM dithiothreitol) and incubated at 30 °C with shaking at 225 rpm. The conditioned yeast cells were then collected via centrifugation at 4600 × g for 3 min at room temperature, washed once with 50 mL ice-cold electroporation buffer and resuspended in electroporation buffer to reach a final volume of 1 mL. The cells were kept on ice until used.

5 μg of the purified linearized vector and 2.7 μg of the purified library insert were added to 400 μL of conditioned yeast. The mixture was transferred to a pre-chilled BioRad GenePulser cuvette with 2 mm electrode gap and kept on ice for 5 min prior to electroporation. Cells were electroporated at 2.5 kV and 25 μF, achieving a time constant between 3.7 and 4.1 ms. Electroporated cells were recovered into 4 mL of YPD media supplemented with 4 mL of 1 M sorbitol and incubated at 30 °C with shaking at 225 rpm for 1 h. Recovered cells were collected via centrifugation at 1700 × g for 3 min at room temperature, resuspended in 0.6 mL SD-CAA medium (2% w/v D-glucose, 0.67% w/v yeast nitrogen base with ammonium sulfate, 0.5% w/v casamino acids, 0.54% w/v Na_2_HPO_4_, and 0.86% w/v NaH_2_PO_4_·H_2_O, all dissolved in deionized water), plated across 3× 150 mm SD-CAA plates (2% w/v D-glucose, 0.67% w/v yeast nitrogen base with ammonium sulfate, 0.5% w/v casamino acids, 0.54% w/v Na_2_HPO_4_, 0.86% w/v NaH_2_PO_4_·H_2_O, 18.2% w/v sorbitol, and 1.5% w/v agar, all dissolved in deionized water) and incubated at 30 °C for 48 h. After 48 h, approximately 200,000 colonies were collected in SD-CAA medium, centrifuged at 4600 × g for 5 min at room temperature, and resuspended in SD-CAA medium with 15% v/v glycerol such that OD_600_ was 50. Glycerol stocks were stored at −80 °C for later use. This protocol was modified from a previously described protocol^49^.

### PacBio sequencing of the combination variant library

Plasmids from the transformed yeast cells were extracted using a Zymoprep Yeast Plasmid Miniprep II Kit (Zymo Research) following the manufacturer’s protocol. The library insert and barcode were amplified using primers PacBio Recovery_F and PacBio Recovery_R (**Table S3**). This PCR was performed using Q5 high-fidelity DNA polymerase (NEB) with the following settings: 98 °C for 30 s, 25 cycles of (98 °C for 10 s, 62 °C for 15 s, 72 °C for 30 s), 72 °C for 2 min, 4 °C indefinitely. PCR products were gel purified and sequenced on one SMRT Cell 8M on a PacBio Sequel IIe using the CCS sequencing mode and a 15-hour movie time.

### Analysis of PacBio sequencing data

Circular consensus sequences (CCSs) were generated from the raw subreads using the ccs program (https://github.com/PacificBiosciences/ccs, version 6.4.0), setting the parameters to require 99.9% accuracy and a minimum of 3 passes. CSSs in FASTQ format were parsed using SeqIO module in BioPython^50^. For each read, the positions of the stem helix peptide, S2P6 heavy and light chain variable domains, as well as the 16-nucleotide barcode were identified by aligning their flanking sequences to the read. If the lengths of these regions deviated from the expected lengths, the read would be discarded. The nucleotide sequences of the stem helix peptide, as well as S2P6 heavy and light chain variable domains were then translated. If the amino acid sequence of the stem helix peptide did not match any of the 27 stem helix peptides, the read would be discarded. Similarly, if the amino acid sequences at the mutated positions of S2P6 did not match our library design, the read would be discarded. Subsequently, the combination variant and the 16-nucleotide barcode in each read were identified.

Since reads that shared the same 16-nucleotide barcode should have the same combination variant, we compared the amino acid sequences at the positions of interest among different reads that shared the same 16-nucleotide barcode. For each position of interest, an amino acid variant present in at least 80% of the reads that shared the same 16-nucleotide barcode was regarded as the true variant. If none of the amino acid variant was present in at least 80% of the reads that shared the same 16-nucleotide barcode, all reads that shared the given 16-nucleotide barcode would be discarded. Besides, those 16-nucleotide barcodes that appeared in only one read were discarded.

### Fluorescence-activated cell sorting (FACS) of yeast display library

A 150 μL glycerol stock of the transformed yeast display library was recovered in 50 mL SD-CAA medium by incubating at 27 °C with shaking at 250 rpm until OD_600_ reached between 1.5-2.0. Then 15 mL of the yeast culture was harvested via centrifugation at 4600 × g at 4 °C for 5 min. The supernatant was discarded, and SGR-CAA induction media (2% w/v galactose, 2% w/v raffinose, 0.1% w/v D-glucose, 0.67% w/v yeast nitrogen base with ammonium sulfate, 0.5% w/v casamino acids, 0.54% w/v Na_2_HPO_4_, and 0.86% w/v NaH_2_PO_4_·H_2_O, all dissolved in deionized water) was added to a final volume of 50 mL with an initial OD_600_ of 0.5. The SGR-CAA yeast culture was transferred to a baffled flask and incubated at 18 °C with shaking at 250 rpm for about 24 h until the OD_600_ reached between 1.3-1.6.

For each staining replicate, 1 mL of yeast culture was harvested via centrifugation at 4600 × g at 4 °C for 5 min. The pellet was washed twice with 1 mL of 1× PBS and resuspended in 1 mL of 1× PBS. For the library binding sort, anti-HA-tag mouse antibody with Alexa Fluor 647 conjugate (6E2, Cell Signaling Technology, Cat. No. 3444) was added to washed cells at a 1:100 dilution. A no-stain negative control was included in which nothing was added to the PBS-resuspended cells. Samples were incubated overnight at 4 °C with rotation. For protease treatment, the stained cells were spun down and the supernatant was removed. 10 μL HRV 3C protease (ThermoFischer, Cat. No. 88946), 10 μL protease reaction buffer, and 80 μL water was added to the stained cells. The cells incubated with protease at 4 °C for 1 h.

The yeast cells were pelleted and washed twice in 1× PBS and resuspended in FACS tubes containing 2 mL 1× PBS. Using a BD FACS Aria II cell sorter (BD Biosciences) and FACS Diva software v8.0.1 (BD Biosciences), cells in the selected gates were collected in 1 mL of SD-CAA containing 1× penicillin/streptomycin. Single yeast cells were gated by forward scatter (FSC) and side scatter (SSC). For the binding sort, single cells were gated into 4 bins along the Alexa Fluor 647 axis. Cells expressing the highest Alexa Fluor 647 fluorescence were sorted into “bin 4”, then the next highest into “bin 3”, followed by “bin 2” and then “bin 1”. Gating strategy used is shown in **Fig. S4A**. Number of cells collected per bin per replicate is shown in **Fig. S4B**. Cells were then collected and grown overnight in SD-CAA at 30 °C with shaking at 225 rpm. FlowJo v10.8 software (BD Life Sciences) was used to analyze FACS data. Replicates were performed on the same day, starting from separate glycerol stocks of the transformed library.

### Illumina sequencing of the post-sorted yeast display library

Plasmids from the sorted yeast cells were extracted using a Zymoprep Yeast Plasmid Miniprep II Kit (Zymo Research) following the manufacturer’s protocol. The 16-nucleotide barcode was amplified using primers Barcode_Recovery_F and Barcode_Recovery_R (**Table S3**), which contained part of the adapter sequence required for Illumina sequencing. A maximum of 100 ng of genomic DNA per 50 μL PCR reaction was used as template. This PCR was performed using Q5 high-fidelity DNA polymerase (NEB) with the following settings: 98 °C for 30 s, 25 cycles of (98 °C for 10 s, 62 °C for 15 s, 72 °C for 15 s), 72 °C for 2 min, 4 °C indefinitely. The PCR products were gel purified. For each sample, 20 ng of the purified PCR product was appended with the rest of adapter sequence and index via PCR using primers: 5’-AAT GAT ACG GCG ACC ACC GAG ATC TAC ACX XXX XXX XAC ACT CTT TCC CTA CAC GAC GCT-3’, and 5’-CAA GCA GAA GAC GGC ATA CGA GAT XXX XXX XXG TGA CTG GAG TTC AGA CGT GTG CT-3’. Positions annotated by an “X” represented the nucleotides for the index sequence. This PCR was performed by PrimeSTAR Max polymerase (Takara Bio) with the following settings: 98 °C for 30 s, 10 cycles of (98 °C for 10 s, 55 °C for 15 s, 72 °C for 15 s), 72 °C for 2 min, 4 °C indefinitely. Indexed products were gel purified, mixed at equimolar ratios, and submitted for next generation sequencing using Illumina NovaSeq X PE150.

### Analysis of Illumina sequencing data

The Illumina NovaSeq sequencing data were obtained in FASTQ format. Forward and reverse reads of each paired-end read were merged by PEAR^51^. The merged reads were then parsed by SeqIO module in BioPython^50^, followed by primer trimming to remove primer sequences. Reads with incorrect primer sequences or lengths were discarded. Reads that did not match any barcodes from our PacBio data were also discarded. The combination variant, which consisted of an S2P6 variant *i* and a stem helix peptide variant *s,* in each read was then identified using the barcode. The frequency (*F*) of a combination variant within bin *n* at timepoint *t* of replicate *r* was computed as follows:

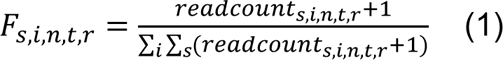

A pseudocount of 1 was added to the read counts of each mutant to avoid division by zero in subsequent steps. The frequency (*F*) of each combination variant among bins at *t* = 0 h (i.e. pre-protease cleavage) was used to calculate the expression score (*E*)^37,52^:

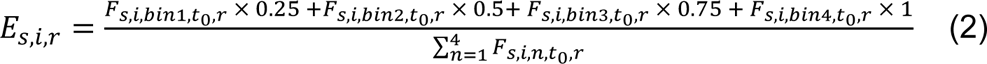

The frequency (*F*) of each combination variant among bins at *t* = 1 h (i.e. post-protease cleavage) was then used to calculate the binding score (*B*):

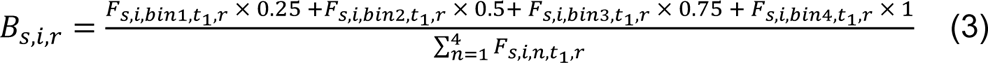

The average binding score for combination variants that contained the negative control peptide (DSAKEALDKYFKNH) were calculated (*B_negctrl_*). The binding score of the combination variant that contained SARS-CoV-2 stem helix peptide (DSFKEELDKYFKNH) and S2P6 WT was also extracted (*B_SARS2-S2P6WT_*). The normalized binding score (*NormB*) of each combination variant *i* was then calculated as followed:

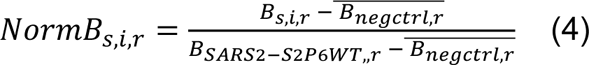

The reported *NormB* was the average between replicates.

### Expression and purification of Fabs

Codon-optimized oligonucleotides encoding S2P6 V_H_ variants were synthesized as eBlocks (Integrated DNA Technologies), and then cloned into phCMV3 vector in Fab format using Gibson assembly. The plasmids were co-transfected into Expi293F cells at a 2:1 (HC:LC) mass ratio using ExpiFectamine 293 Reagent (Thermo Fisher Scientific) following the manufacturer’s protocol for a 25 mL culture. 6 days post-transfection, the supernatant of each Expi293F culture was collected and clarified by centrifugation at 4600 × g for 45 min at 4 °C to remove cells and debris. After clarification, the supernatant was filtered through a 0.22 μm polyethersulfone membrane filter (Millipore).

Fabs were purified using IgG-CH1-XL beads (Thermo Fisher Scientific). The beads were washed with MilliQ H_2_O three times and resuspended in 1× PBS. The clarified and filtered supernatant was incubated with washed beads overnight at 4 °C with gentle rocking. The supernatant with beads was then loaded into a column. The flowthrough was discarded, and the beads were washed once with 1× PBS. To elute the antibody, the beads were incubated in 60 mM sodium acetate, pH 3.7 for 10 min at 4 °C and the flow-through was collected. The eluate containing Fab was buffer exchanged into 1× PBS using a centrifugal filter unit with a 10 kDa molecular weight cut-off (Millipore). Fabs were stored at 4 °C.

### Biolayer interferometry binding assays

Biolayer interferometry (BLI) experiments were performed at room temperature using Octet Red96e instrument (Sartorius). Biotin-labeled stem peptides (GenScript) at 4 μg/mL in 1× kinetics buffer (1× PBS, pH 7.4, 0.01% w/v BSA, and 0.002% v/v Tween 20) were loaded onto streptavidin (SA) biosensors and incubated with Fabs at 500 nM, 750 nM, and 1000 nM. The assay consisted of five steps: (1) baseline: 60 s with 1× kinetics buffer; (2) loading: 120 s with biotin-labeled stem peptides; (3) baseline: 60 s with 1× kinetics buffer; (4) association: 120 s with Fab samples; and (5) dissociation: 120 s with 1× kinetics buffer. For estimating the exact K_D_, a 1:1 binding model was used. When a 1:1 binding model did not fit well due to non-specific binding, a 2:1 heterogeneous ligand model was used to improve fitting.

## DATA AVAILABILITY

Raw deep sequencing data generated in this study have been submitted to the NIH Sequence Read Archive under accession number: PRJNA1064076 and PRJNA1113356.

## CODE AVAILABILITY

Custom codes for data analysis have been deposited to: https://github.com/nicwulab/S2P6_lib-on-lib_screen.

## ACKNOWLEDGMENTS

We thank the Roy J. Carver Biotechnology Center at the University of Illinois Urbana-Champaign for assistance with fluorescence-activated cell sorting and performing next-generation sequencing. This work was supported by the US National Institutes of Health DP2 AT011966 (N.C.W.), R01 AI167910 (N.C.W.), Searle Scholars Program (N.C.W.), National Institutes of the Health Chemical Biology Interface Training Program T32 GM136629 (M.Y.O.), and National Science Foundation Graduate Research Fellowship DGE 21 46756 (M.Y.O.).

## AUTHOR CONTRIBUTIONS

M.Y.O. and N.C.W. designed research. M.Y.O. performed research. M.Y.O., W.O.O., and N.C.W. analyzed data. M.Y.O. and N.C.W. wrote the paper.

## COMPETING INTERESTS

N.C.W. serves as a consultant for HeliXon. All authors declare no other competing interests.

