## Supplemental Information for "A library-on-library screen reveals the breadth expansion landscape of a broadly neutralizing betacoronavirus antibody"

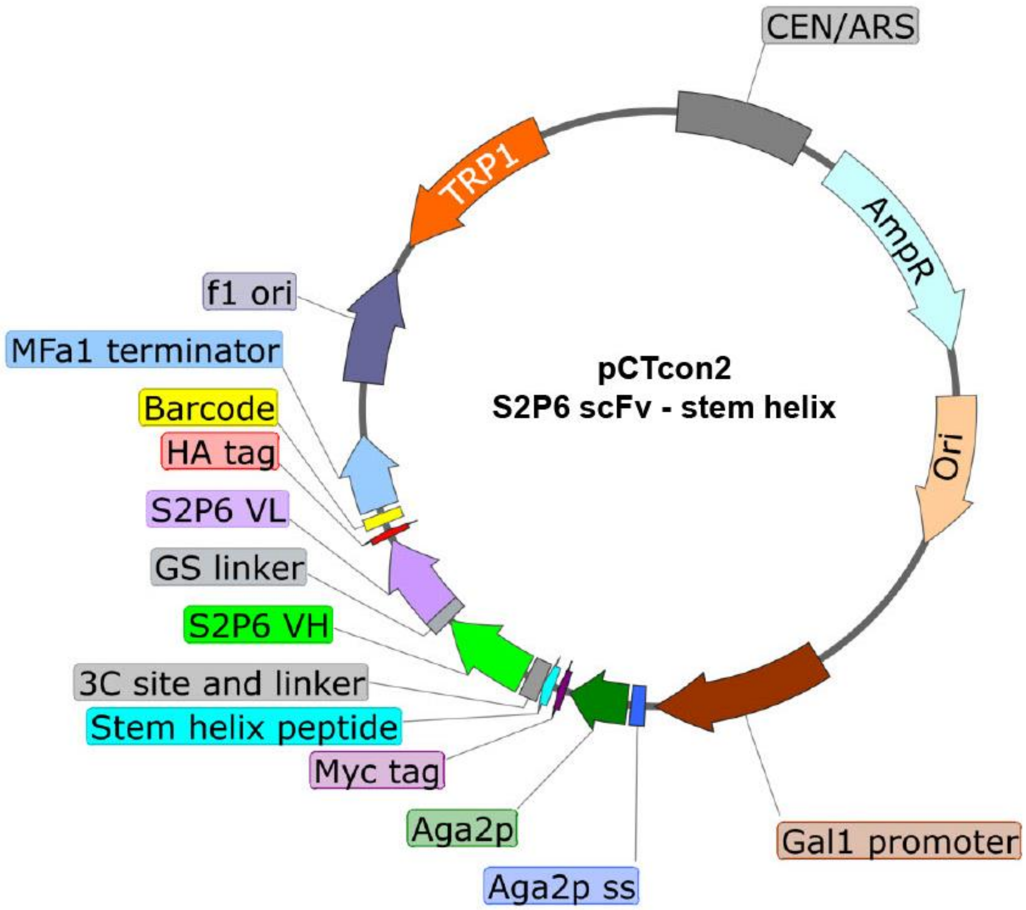


**Figure S1. Plasmid map details.** Schematic of our plasmid construct for yeast display library-on-library screen with key features highlighted.

**A**

```
          1                              32              48       55 56
UCA_VH   QVQLVQSGAEVKKPGASVKVSCKASGYTFTSYYMHWVRQAPGQGLEWMGIINPSGGSTSY
S2P6_VH  EVQLVQSGAEVKKPGASVKVSCKASGYTFTSQYMHWVRQAPGQGLEWIGIINPSGVHTSY

                   69       78
UCA_VH   AQKFQGRVTMTRDTSTSTVYMELSSLRSEDTAVYYCARGSPKGAFDYWGQGTLVTVSS
S2P6_VH  AQKFQGRVTLTRDTSTSTLYMELSSLRSEDTAVYYCARGSPKGAFDYWGQGTLVTVSS
```

**B**

```
            4 5                        29 31
UCA_VL   EIVLTQSPGTLSLSPGERATLSCRASQSVSSSYLAWYQQKPGQAPRLLIYGASSRATGIP
S2P6_VL  EIVMMQSPGTLSLSPGERATLSCRASQSVRSNYLAWYQQKPGQAPRLLIYGASSRATGIP

                                                    105
UCA_VL   DRFSGSGSGTDFTLTISRLEPEDFAVYYCQQYGSSPPRFTFGPGTKVDIK
S2P6_VL  DRFSGSGSGTDFTLTISRLEPEDFAVYYCQQYGSSPPRFTFGPGTKVEIK
```

**C**

| scFv residue | Variants in library |
|---|---|
| $V_H$ 32 | Y, Q |
| $V_H$ 48 | M, I |
| $V_H$ 55 | G, V |
| $V_H$ 56 | H, S |
| $V_H$ 69 | M, L |
| $V_H$ 78 | V, L |
| $V_L$ 4 | L, M |
| $V_L$ 5 | T, M |
| $V_L$ 29 | S, R |
| $V_L$ 31 | S, N |

**Figure S2. S2P6 variant library design. (A-B)** Sequence alignments of **(A)** S2P6 $V_H$ and UCA, as well as **(B)** S2P6 $V_L$ and UCA. Somatic hypermutations are highlighted in red. **(C)** Amino acid variants in the S2P6 variant library.

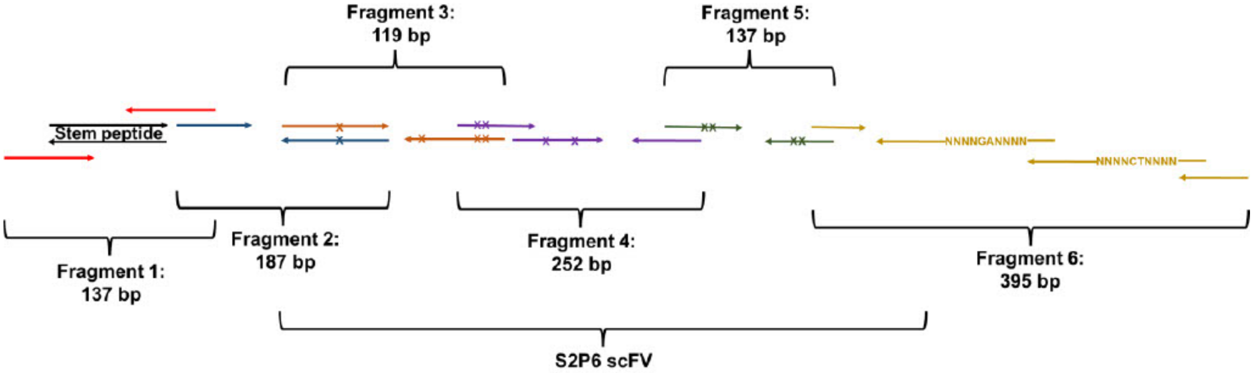


**Figure S3. Strategy for assembling the combination variant library.** The combination variant library was assembled via overlap PCR of six fragments (represented by the color of primers) encoding (from 5' to 3') the stem helix peptide variant library, S2P6 variant library, linkers, and 16-nucleotide barcode (NNNNGANNNN … NNNNCTNNNN).

**A**

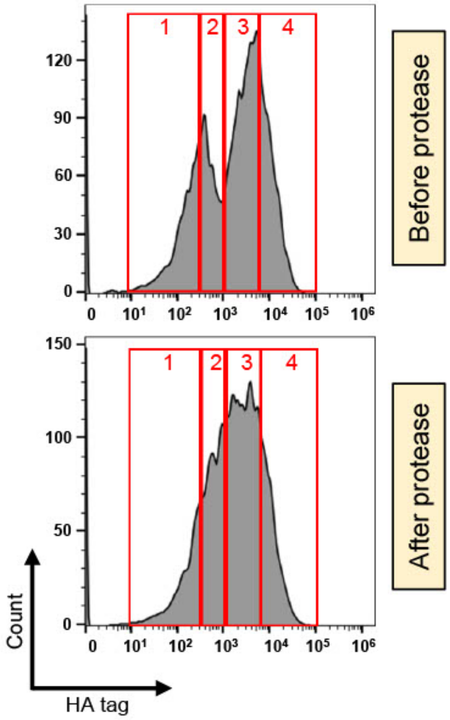


**B**

| Protease | Replicate | Gate | Cell count |
|---|---|---|---|
| Before | 1 | 1 | 319,217 |
| Before | 1 | 2 | 317,819 |
| Before | 1 | 3 | 554,411 |
| Before | 1 | 4 | 403,008 |
| Before | 2 | 1 | 318,096 |
| Before | 2 | 2 | 324,247 |
| Before | 2 | 3 | 573,971 |
| Before | 2 | 4 | 472,848 |
| After | 1 | 1 | 351,057 |
| After | 1 | 2 | 847,861 |
| After | 1 | 3 | 1,453,835 |
| After | 1 | 4 | 563,505 |
| After | 2 | 1 | 321,741 |
| After | 2 | 2 | 795,598 |
| After | 2 | 3 | 1,694,468 |
| After | 2 | 4 | 845,198 |

**Figure S4. Cell sorting of the combination variant library. (A)** Gating scheme for cell sorting of the combination variant library. Four gates were used (i.e. 1, 2, 3, 4) where 1 represents lowest HA tag signal (Alexa Flour 647 nm) and 4 represents highest HA tag signal. Four library samples were sorted in total: two before protease cleavage and two after protease cleavage. **(B)** Cell counts listed for each sample.

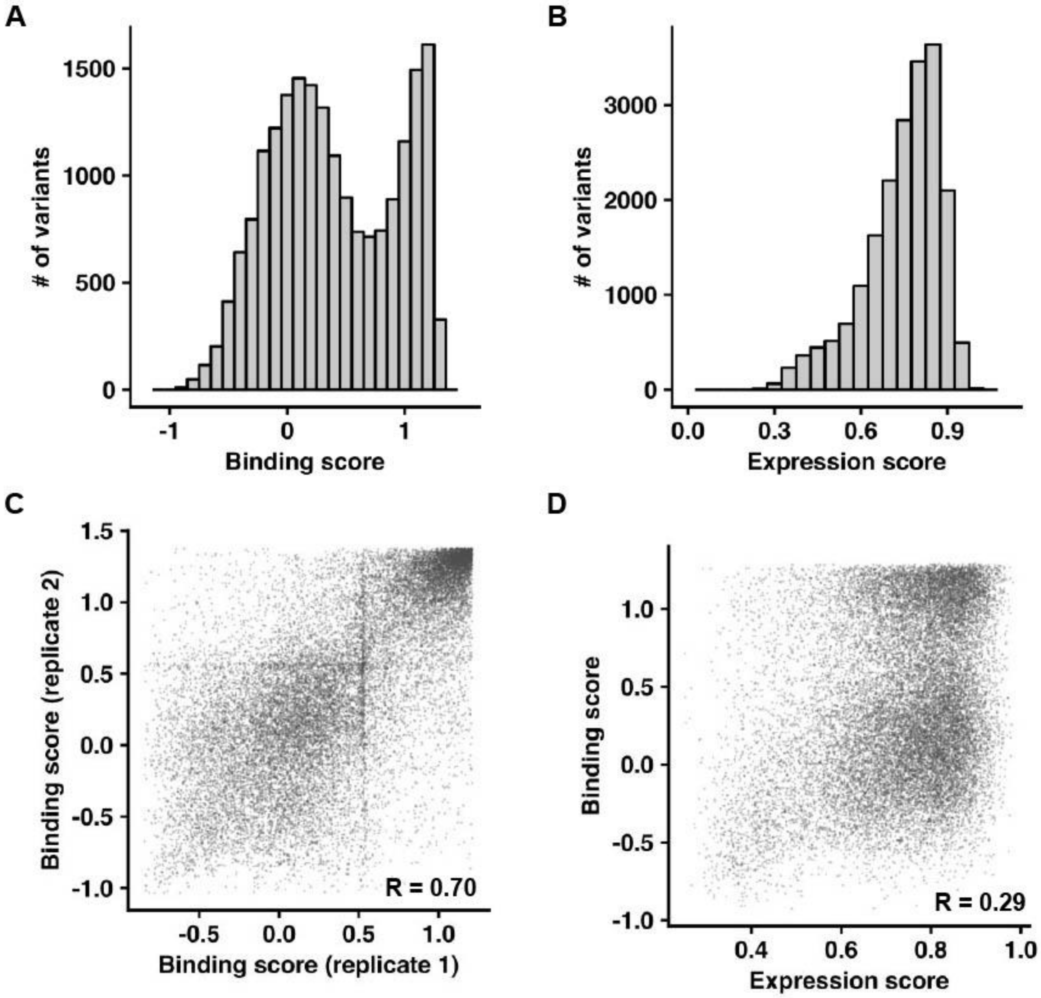


**Figure S5. Expression and binding scores of individual variants in the combination variant library. (A)** Binding score distribution. **(B)** Expression score distribution. **(C)** Comparison of binding scores between two experimental replicates. **(D)** The relationship between binding score and expression score.

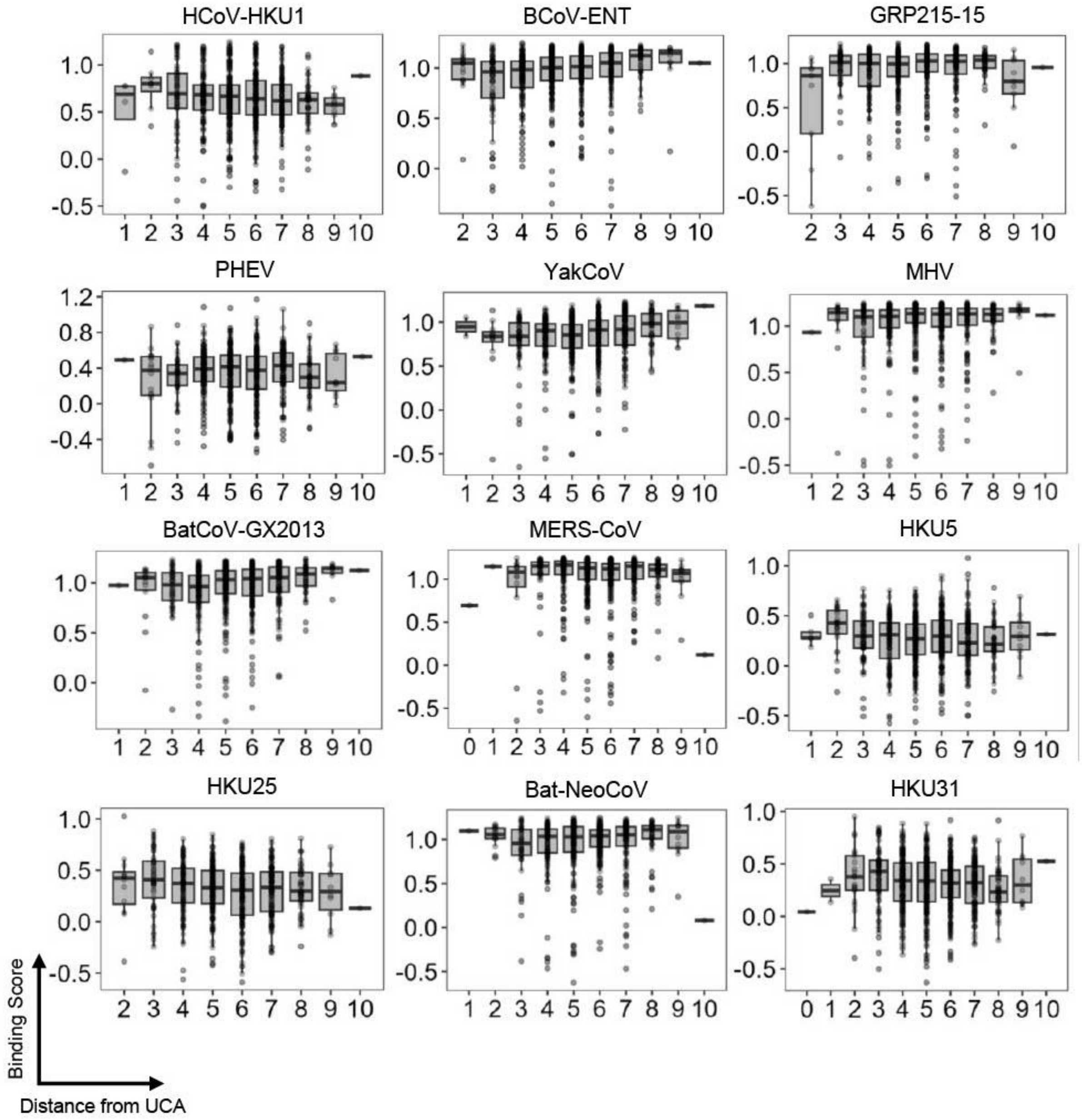
HCoV-HKU1
BCoV-ENT
GRP215-15
PHEV
YakCoV
MHV
BatCoV-GX2013
MERS-CoV
HKU5
HKU25
Bat-NeoCoV
HKU31
Binding Score
Distance from UCA

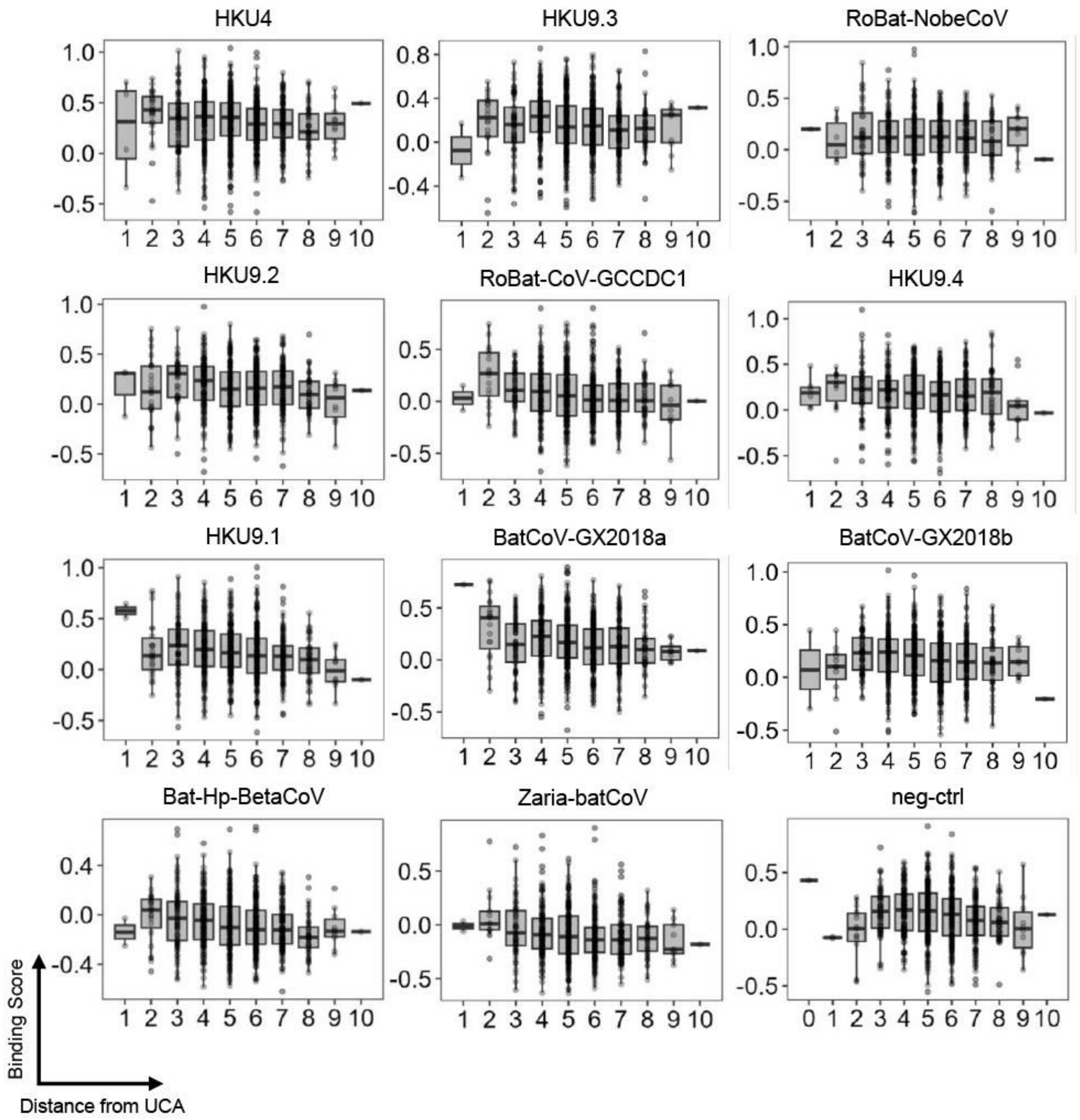


**Figure S6. Relationship between binding score and distance from UCA.** Binding scores of S2P6 variants with different numbers of somatic hypermutations (i.e. distance from UCA) to the indicated stem helix peptides are shown. Each data point represents the one S2P6 variant. For the boxplot, the middle horizontal line represents the median. The lower and upper hinges represent the first and third quartiles, respectively. The upper whisker extends to the highest data point within a 1.5× inter-quartile range (IQR) of the third quartile, whereas the lower whisker extends to the lowest data point within a 1.5× IQR of the first quartile.

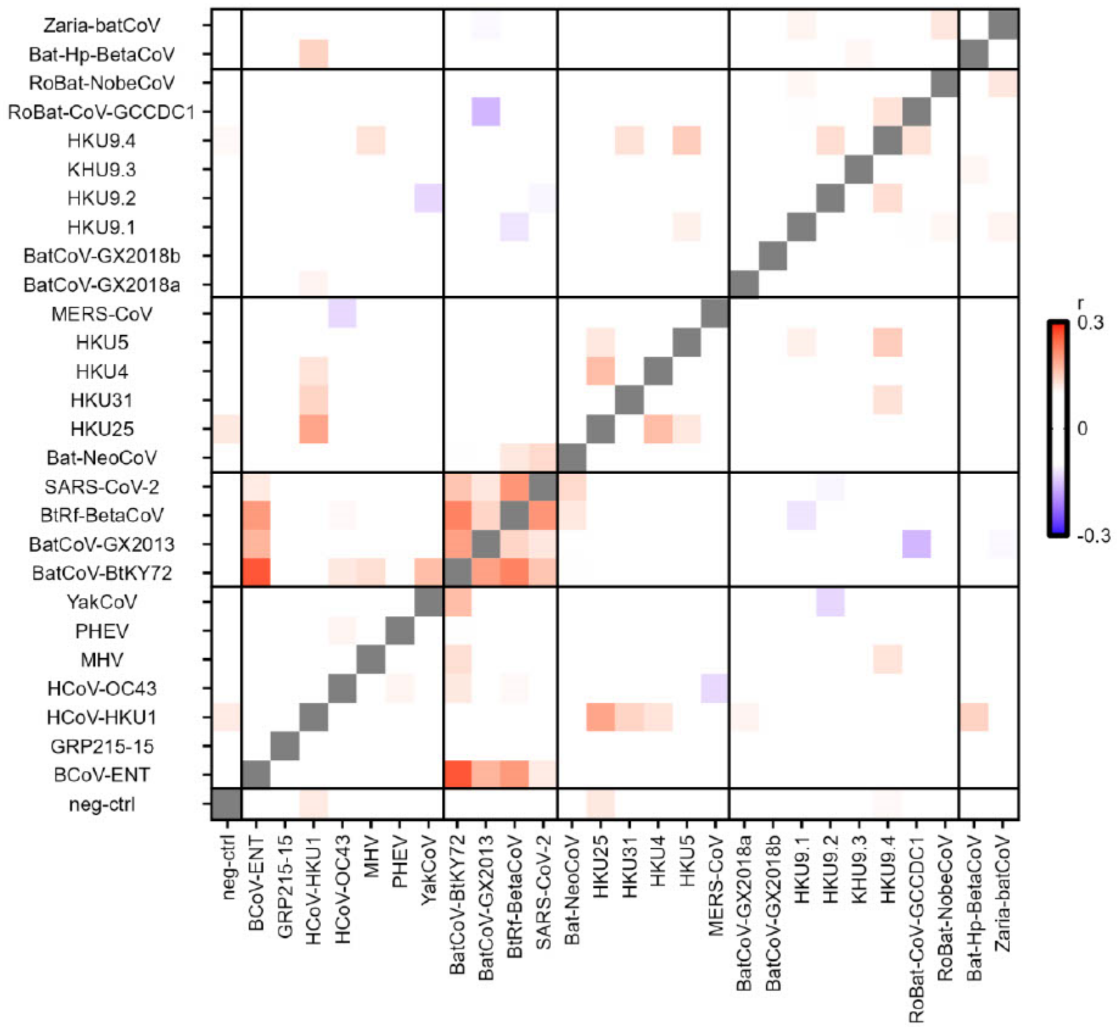


**Figure S7. Correlation of binding scores between stem helix peptides.** Pairwise Pearson correlations of binding scores between stem helix peptides. Red indicates positive correlation coefficient, whereas blue indicates negative correlation coefficient.

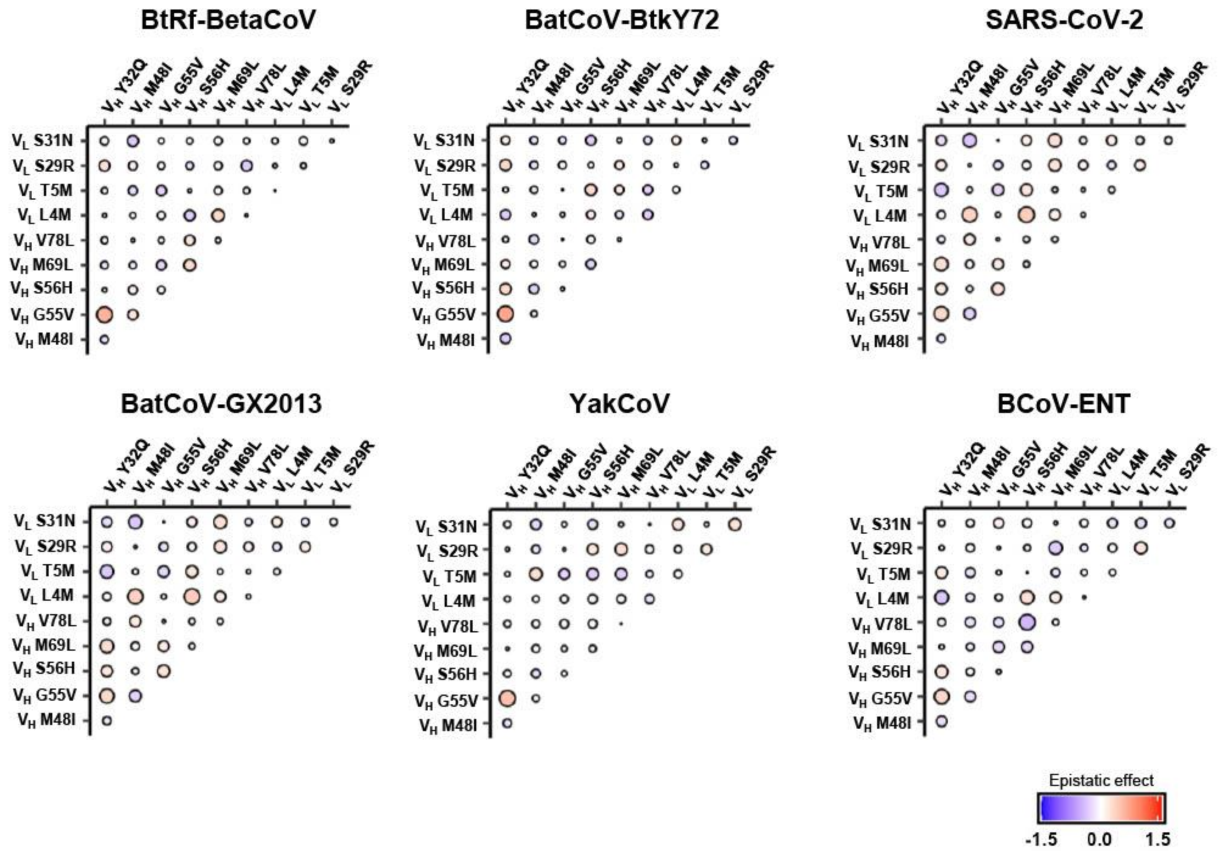


**Figure S8. Pairwise epistasis between S2P6 somatic hypermutations.** Inference of epistatic interactions between S2P6 somatic hypermutations for binding to BtRf-BetaCoV, BatCoV-BtkY72, SARS-CoV-2, BatCoV-GX2013, YakCoV, and BCoV-ENT. Red and blue represents positive and negative epistasis, respectively. Magnitude is proportional to the size of the circle.

### BatCoV-BtkY72

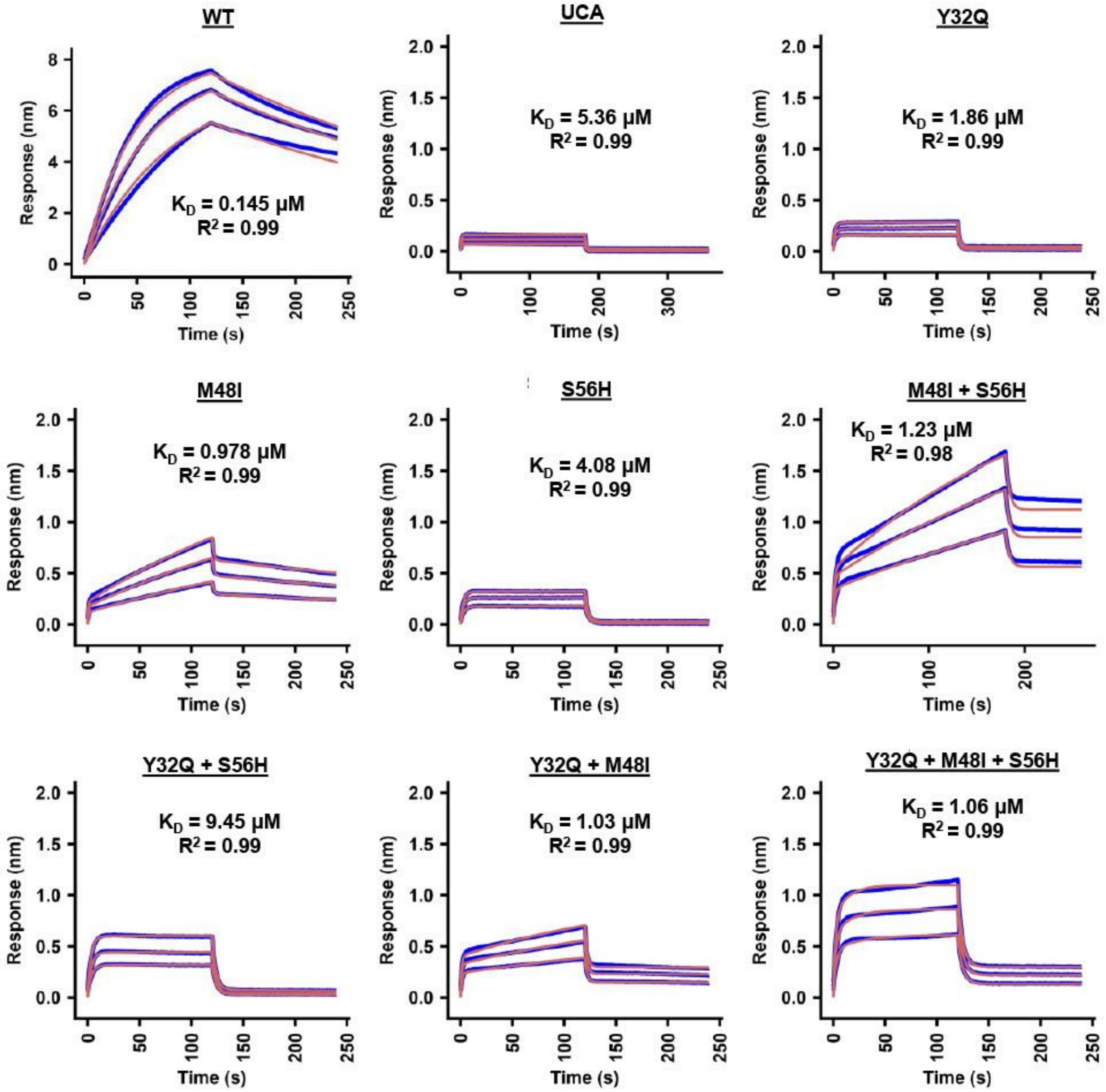

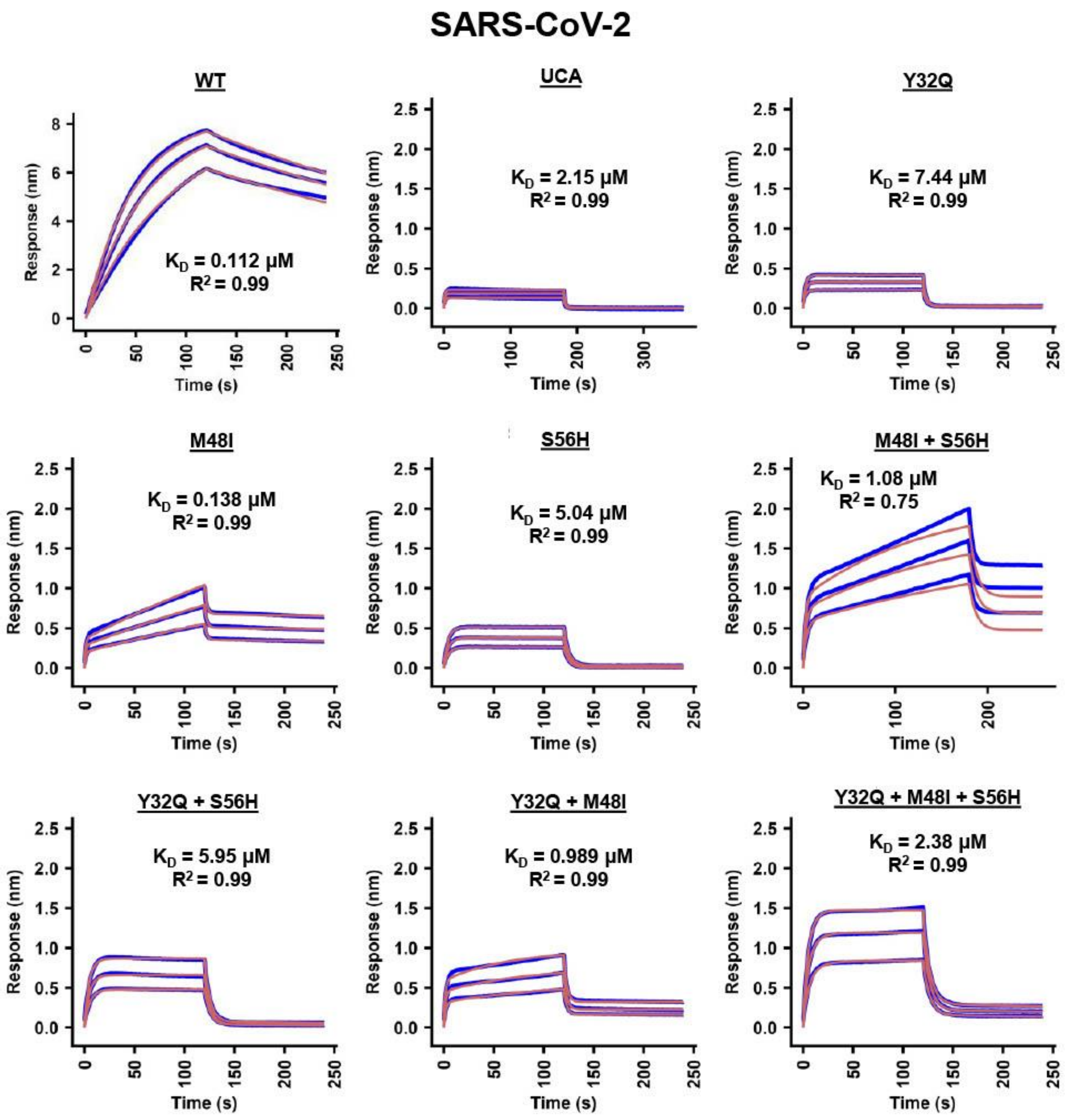

SARS-CoV-2
WT
$K_D$ = 0.112 µM
$R^2$ = 0.99
UCA
$K_D$ = 2.15 µM
$R^2$ = 0.99
Y32Q
$K_D$ = 7.44 µM
$R^2$ = 0.99
M48I
$K_D$ = 0.138 µM
$R^2$ = 0.99
S56H
$K_D$ = 5.04 µM
$R^2$ = 0.99
M48I + S56H
$K_D$ = 1.08 µM
$R^2$ = 0.75
Y32Q + S56H
$K_D$ = 5.95 µM
$R^2$ = 0.99
Y32Q + M48I
$K_D$ = 0.989 µM
$R^2$ = 0.99
Y32Q + M48I + S56H
$K_D$ = 2.38 µM
$R^2$ = 0.99
Response (nm)
Time (s)

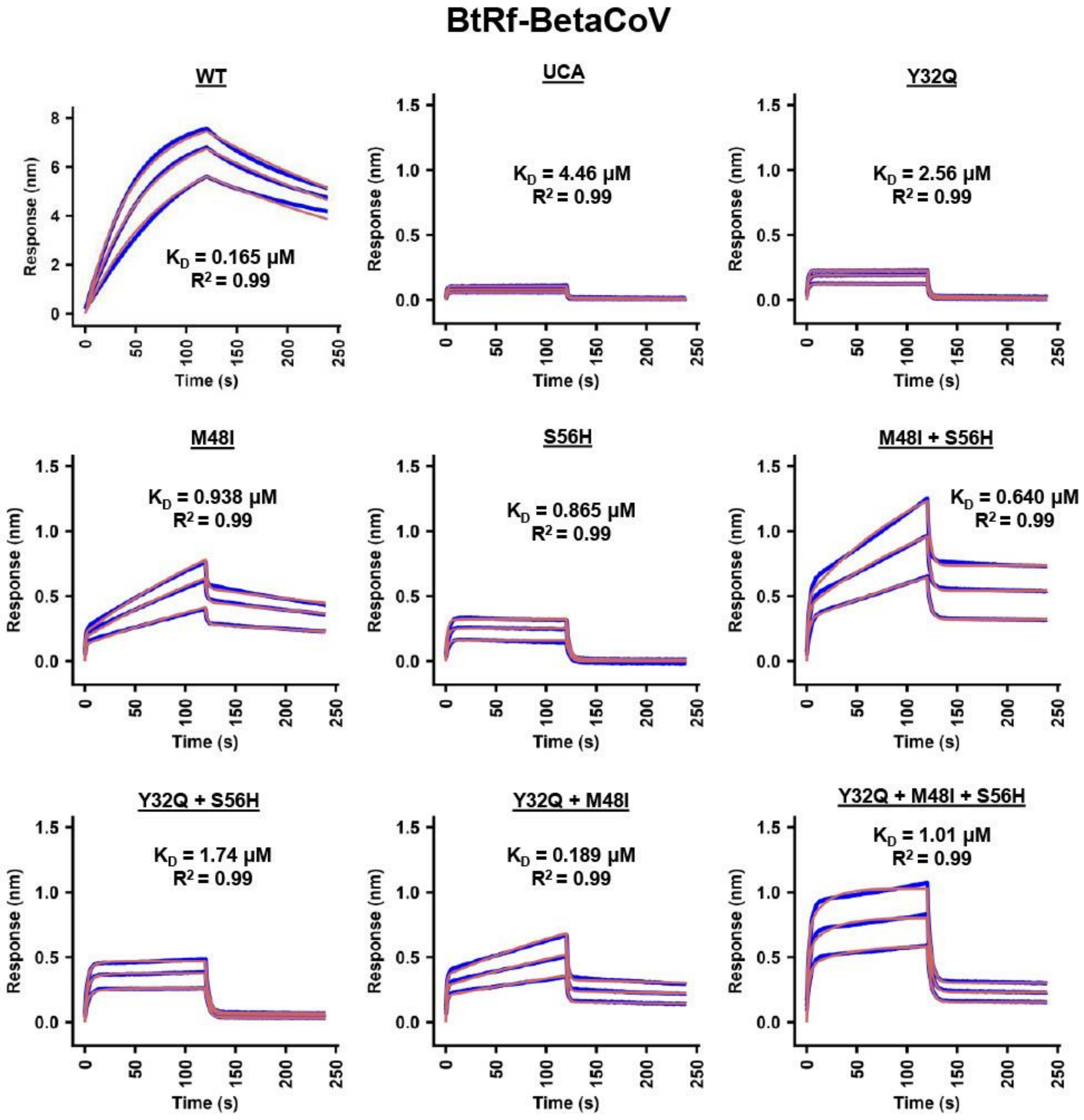
BtRf-BetaCoV
WT
K_D = 0.165 μM
R² = 0.99
UCA
K_D = 4.46 μM
R² = 0.99
Y32Q
K_D = 2.56 μM
R² = 0.99
M48I
K_D = 0.938 μM
R² = 0.99
S56H
K_D = 0.865 μM
R² = 0.99
M48I + S56H
K_D = 0.640 μM
R² = 0.99
Y32Q + S56H
K_D = 1.74 μM
R² = 0.99
Y32Q + M48I
K_D = 0.189 μM
R² = 0.99
Y32Q + M48I + S56H
K_D = 1.01 μM
R² = 0.99
Response (nm)
Time (s)

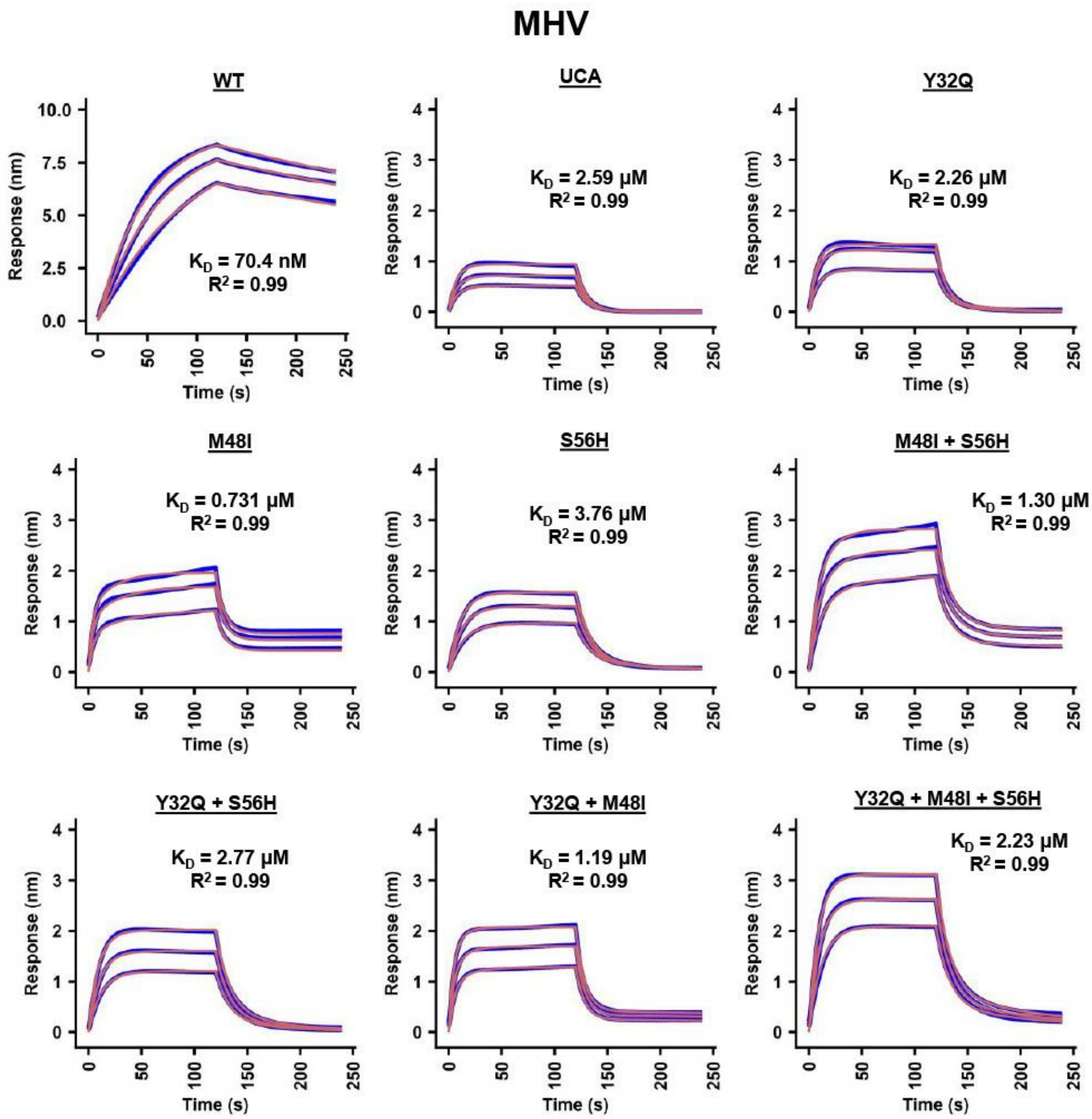


**Figure S9. Binding kinetics of recombinantly expressed S2P6 variants.** Biolayer interferometry sensorgrams of Fabs (S2P6 WT, UCA, and UCA mutants) binding to BatCoV-BtkY72, SARS-CoV-2, BtRf-BetaCoV, MHV, and HCoV-HKU1 stem helix peptides. Blue lines represent response curve and red lines represent best fit model (1:1 binding model or 2:1 heterogeneous ligand model). Binding kinetics were measured for three Fab concentrations (500 nM, 750 nM, and 1000 nM). Dissociation constant ($K_D$) and the goodness of model fitting ($R^2$) are included on each plot.
